# A pseudo-phased genome assembly for *Hemileia vastatrix* reveals an isolate-specific chromosomal haploid trisomy

**DOI:** 10.64898/2026.04.04.716458

**Authors:** Peri A. Tobias, Richard J. Edwards, Jamie Botting, Giullia di Lorenzo, Vera Inácio, Inês Diniz, Maria do Céu Silva, Leonor Guerra-Guimarães, Vitor Várzea, Robert F. Park, Dora Batista

**Author notes:** Instituto de Biosistemas e Ciências Integrativas (BioISI), Faculdade de Ciências, Universidade de Lisboa, 1749-016 Lisboa, Portugal (Vera Inácio). National Institute for Agricultural and Veterinary Research (INIAV, I.P.), Estrada de Leiria, 2460-059 Alcobaça, Portugal. equal first authors (Peri A. Tobias, Richard J. Edwards).

## Abstract

Recurrent epidemics of coffee leaf rust, caused by the fungal pathogen *Hemileia vastatrix,* have constrained production of Arabica coffee for over 150 years. Here, we present a pseudo-phased, chromosome-level genome resource for *H. vastatrix,* isolate Hv178a, to guide research into disease management. The Hv178a genome assembly is 665 and 638 Mbp for haplotype A and B respectively, localised to 18 chromosomes. We determined that the genomes are highly repetitive at ∼90%, with a GC content of ∼33%. We present the full annotation of 13,760 and 17,998 protein coding genes, and we predicted 452 and 496 effectors in haplotype A and B respectively. Depth-based comparisons with 11 additional *H. vastatrix* isolates revealed increased chromosome 17 (chr17) copy number in Hv178a. Validation with qPCR supports a chr17 trisomy in Hv178a absent from the ancestral lineage and potentially explaining the observed change in virulence.

## Introduction

The specialized fungal pathogen *Hemileia vastatrix* Berk. & Broome (phylum Basidiomycota, class Pucciniomycetes, order Pucciniales), causing the devastating disease coffee leaf rust (CLR), has been the major constraint to Arabica coffee (*Coffea arabica*) production for more than one and a half centuries (Cabral et al., 2016; McCook and Vandermeer, 2015; Talhinhas et al., 2017). First recorded by an English explorer in 1861 near Lake Victoria (East Africa) on wild *Coffea* species, CLR wiped out Arabica coffee cultivation from Ceylon (Sri Lanka) only five years later, with devastating social and economic consequences (Morris, 1880). Since this historical first outbreak, the disease gained a worldwide distribution, spreading progressively through coffee production areas of Asia and Africa, and, finally, crossing the Atlantic Ocean where it expanded across Latin America (McCook, 2006). With the more recent detection of CLR in Hawaii in 2020 (Keith et al., 2021), the disease became endemic in all coffee producing regions of the world. Although the development and cultivation of rust resistant varieties have contributed to CLR control, high adaptation of the fungus has led to recurrent and severe epidemics resulting in heavy yield and revenue losses. Reports estimate production losses more than $1 billion annually worldwide (Kahn, 2019).

Due to the ability of the pathogen to respond rapidly to selection pressure, more than 55 physiological races have been identified to date (Silva et al., 2022). The rapid evolution of *H. vastatrix* to overcome resistance in coffee cultivars is more puzzling considering that the pathogen exists almost exclusively in the asexual stage of its life cycle, and that the dikaryotic urediniospores are the only known functional propagules (Koutouleas et al., 2019; Silva et al., 2018; Silva et al., 2022; Talhinhas et al., 2017). The isolate used for the current research is a mutant virulent strain obtained in 1960 from the greenhouse complex of Centro de Investigação das Ferrugens do Cafeeiro (CIFC), named CIFC Hv178a (Figure 1).

**Figure 1.**
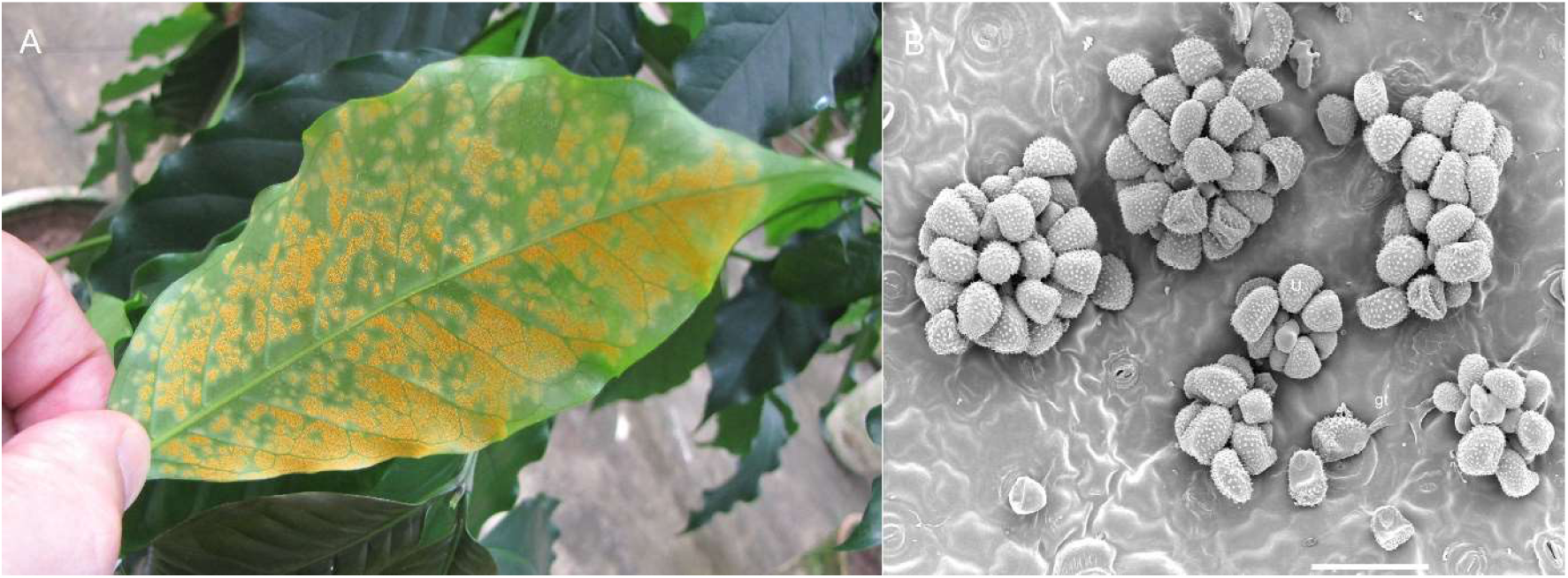
A *Coffea arabica* leaf with symptoms of coffee leaf rust disease caused by the fungal pathogen *Hemileia vastatrix* at The Centro de Investigação das Ferrugens do Cafeeiro (CIFC) greenhouse complex, Universidade de Lisboa, Portugal in 2022. B Surface view of superstomatal uredosporic sori of H. vastatrix. Note the reniform uredospores (U), and an early stage of germination (gt = germ tube). Scanning Electron Microscopy (Bar = 50 µm) (HG Azinheira and MC Silva, unpublished.)

Deconstructing the evolutionary complexity between *H. vastatrix* and its only known host is crucial to developing options for disease control. Such goals rely on the availability of a comprehensive and high-quality genome reference assembly. Here, using additional chromosome confirmation capture sequence data, we manually curated to improve on a previous haploid genome assembly (Tobias et al. 2022) and report a pseudo-phased and chromosome-level genome for *H. vastatrix* strain CIFC Hv178a, henceforth Hv178a. We confirm the conservation of 18 haploid chromosomes within *H. vastatrix*, in line with numbers determined for distantly related rust fungi (Luo et al., 2024; Schwessinger et al., 2022). We compared the assembled genome to data from 11 other strains, including the ancestral Hv178, and determined intriguing chromosome copy number variation indicating trisomy for chromosome 17, validated with qPCR, potentially explaining the observed change in virulence. The genome provides a high-quality resource for advancing knowledge on the complex coffee rust pathogen as well as identifying a novel chromosomal anomaly for this isolate.

## Methods

### Fungal material

*Hemileia vastatrix* isolate Hv178a (type specimen of race XIV: virulence profile v_2,3,4,5_), maintained at the spore collection of Centro de Investigação das Ferrugens do Cafeeiro (CIFC), Instituto Superior de Agronomia, Universidade de Lisboa, was used in this work. This isolate is a mutant strain obtained in 1960 at CIFĆs greenhouse complex derived from isolate CIFC Hv178 (race VIII: virulence profile v_2_,_3,5_), collected in India (Balehonnur, Coffee Research Station) in 1958 from *Coffea arabica* S.288-23. Within the scope of experimental selection assays for increased virulence, the repeated inoculation of Hv178 onto a resistant Arabica coffee plant led to the emergence of a new isolate, Hv178a, bearing the addition of v4 to its virulence spectrum (Bettencourt, 1981). Hv178a was multiplied on its differential host plant (*C. arabica* accession CIFC H147/1, carrying the resistance factors S_H_2,3,4,5), as previously described (d’ Oliveira, 1954). Single uredospore pustules were collected, frozen in liquid nitrogen, and stored at -80°C until further processing.

### DNA extraction and sequencing

High molecular weight (HMW) genomic DNA was extracted using a cetyltrimethylammonium bromide (CTAB)-based modified protocol (Schwessinger and Rathjen, 2017). DNA samples were further purified using Pacific Biosciences (PacBio) SampleNet – Shared Protocol (https://www.pacb.com/wp-content/uploads/2015/09/SharedProtocol-Extracting-DNA-usinig-Phenol-Chloroform.pdf). DNA concentration and purity were measured using a Multiskan SkyHigh Microplate Spectrophotometer (Thermofisher Scientific) and a Qubit 4 Fluorometer (Thermofisher Scientific), and its quality checked on an agarose gel. Ten micrograms of high-quality HMW DNA were sent to CD Genomics (New York, USA) for SMRT bell size select long read library preparation and PacBio Sequel II sequencing. The same sample of HMW DNA was used for library construction with TruSeq DNA PCR-Free kit, following manufacturer’s recommendations, and resequencing with paired-end 151 bp reads at CD Genomics on Illumina an NovaSeq platform.

### Cross-linking of spores and chromatin conformation capture sequencing

To improve the scaffolding of our genome assembly we obtained chromatin conformation sequence data (HiC). About 400 mg of Hv178a urediniospores were crosslinked in a 1% formaldehyde solution for 20 min at room temperature. Crosslinking was quenched by adding glycine to a final concentration of 125 mM. The fixed tissue was then ground in liquid nitrogen and the stored frozen powder sent to Phase Genomics (Seattle, WA) for nuclei isolation, HiC library preparation and sequencing. Data was generated using a Phase Genomics (Seattle, WA) Proximo Hi-C 4.0 Kit, which is a commercially available version of the HiC protocol (Lieberman-Aiden et al., 2009) using the DPNII restriction enzyme. Sequencing was performed on an Illumina HiSeq 4000, generating a total of 175,294,501 PE 150 read pairs (total reads 350,589,002).

### Draft genome assembly

We received raw PacBio Sequel (Menlo Park, California, USA) SMRT™ continuous long reads (CLR) from CD Genomics (New York, USA). Prior to correction we had 7,539,679 reads (166 Gigabases), with a median read length of 17,992. This provided approximately 200 times coverage on an expected genome size of 800 Mb, based on flow cytometry (Tavares et al., 2014). An earlier version of the Hv178a strain genome assembly using this Sequel data was made public (Tobias et al., 2022). The genome indicated a potential *associated fungal genome* (AFG) or contaminant, and this caused problems when attempting to phase the diploid assemblies due to high levels of duplicated gene models. Additionally, our earlier version was scaffolded with limited HiC data, only 31,666,891 PE reads. We therefore decided to reassemble the genome, curate and scaffold with adequate HiC read data coverage. We used Canu (v2.1.1) (Koren et al., 2016) with the following recommended parameters for diploid genome assembly; genomeSize=800m, corOutCoverage=100, corMaxEvidenceErate=0.15, correctedErrorRate=0.040, batOptions=“-dg 3 -db 3 -dr 1 -ca 500 -cp 50”. These options produced an assembly of 1.71 Gb with 6,088 contigs, an N50 of 74,5640 and L50 of 438. Genome outputs were tested for contiguity with Quast version 5.0.2 (Gurevich et al., 2013), and conserved single copy gene completeness with BUSCO (v3.0) (Simão et al., 2015) with the basidiomycota_odb9 database. We used this output going forward.

### Genome deduplication

We ran Purge Haplotigs (v1.0) (Roach et al., 2018) to deduplicate the 1.7 Gb genome using these counts per read depth; -l 32 -m 115 -h 185 after mapping with Minimap2 (v2.3) (Li, 2018). We used these purge parameters for the cut-offs; -I 250M -a 75 and produced two genome fasta outputs with the following BUSCO statistics; C:93.1%[S:57.3%,D:35.8%],F:2.4%,M:4.5% and C:71.0%[S:49.7%,D:21.3%],F:4.1%,M:24.9% for the primary (575M) and haplotig (785M) assemblies. The BUSCO statistics indicated difficulties in separating the haplotypes and the continued presence of the contaminant AFG sequence, with large numbers of duplicated conserved genes in each assembly.

### Assembly decontamination

The goals of the initial assembly curation were to (1) remove the AFG sequences and any other contamination, and (2) purge excess haplotigs. Due to the complications arising from the Purge Haplotigs run, the original Canu primary (464 contigs, 596.4 Mbp) and alternative (2261 contigs, 814.7 Mbp) assemblies were used as the basis for curation. Sequencing reads were mapped onto the earlier Hv178a strain genome assembly (Tobias et al., 2022), using Minimap2 (v2.2) (Li, 2018) for PacBio and BWA mem (v0.7.17) (Li, 2013) for Illumina reads. Mapped reads were partitioned using Samtools (v1.15) (Li et al., 2009) into (1) AFG reads, (2) mtDNA reads, (3) nuclear reads. PacBio nuclear reads were mapped independently onto the primary and alternative Canu assemblies using Minimap2 (v2.2) (Li, 2018). Any contigs with below 3X mean coverage were removed from the assembly as likely contamination (e.g. AFG) or mtDNA. In addition, Tiara (v1.0) (Karlicki et al., 2022) identified one mitochondrial and two bacterial contigs, which were removed from the assembly. Next, Diploidocus (v1.5.0) (Chen et al., 2022) was run on each haplotype using the filtered nuclear PacBio reads for depth calculations, and the filtered nuclear Illumina reads for kmer analysis. Genome completeness was estimated using BUSCO (v5.1.2) (Manni et al., 2021a) with basidiomycota_odb10 database. Following removal of the low coverage contigs, the haplotypes were 64.4% (pri) and 68.8% (alt) complete with 15.9% and 23.6% duplication. However, the combined diploid assembly was 85.5% complete and 80.6% duplicated, suggesting that many BUSCO-containing contigs had both haplotigs assigned to the same haploid assembly. This was exacerbated by Diploidocus, which over-purged because of these false duplications, reducing the diploid BUSCO completeness to 82.9% complete and 50.1% “duplicated”.

### Assembly phase correction and curation

The Diploidocus cleanup erroneously purged contigs that were misassigned to haplotypes rather than falsely duplicated in the overall assembly. To compensate for this, the quarantined contigs from each haploid assembly were added back to the other assembly (e.g. contigs purged from the pri assembly were added to the alt assembly and vice versa). Haplotypes were renamed as hap1 (83.35% Complete, 9.4% Duplicated) and hap2 (76.1% Complete, 5.4% Duplicated) and each haploid assembly subjected to a second round of Diploidocus tidying. This was more successful but still suffered from an overall loss of diploid BUSCO Complete and Duplicated genes. Contigs were therefore manually filtered using Diploidocus sequence statistics and identifying contigs with common BUSCO genes. Where a quarantined contig contained a BUSCO gene that was found twice in one haplotype but not in the other, it was moved to the other haplotype. Excess copies (e.g. once both haplotypes contained the genes) were marked for removal as usual. Contigs were dealt with in size order, largest first, so the smaller contig was always marked for removal. Next, under-quarantined contigs that contained within-haplotype Duplicated BUSCO genes but had not been marked for removal were processed in a similar fashion, moving one copy when the other haplotype lacked the corresponding BUSCO genes, or marking it for removal when it did. The exception were contigs that also contained single-copy BUSCO genes, or Duplicated BUSCO genes that were rated as True Duplicates based on DepthKopy (Chen et al., 2022) read depths. These contigs were retained in the original haplotype. After phase correction, the two haploid assemblies were 83.6% and 78.3% Complete, with 1.9% Duplicated BUSCOs in each case. The combined diploid was 85.3% Complete and 76.6% Duplicated, indicating that most of the assembly was now partitioned correctly into two haplotypes. It should be noted, however, that some loss and misassignment of contigs is likely to remain. These haploid assemblies were then scaffolded to chromosome-level using the HiC data.

### Chromosome-scale scaffolding with HiC reads

Phased genome outputs were then independently scaffolded by using the Aiden Lab pipelines (https://github.com/aidenlab). Firstly, the files were run through the Juicer pipeline (v1.6) (Durand et al., 2016) with default parameters. The final output from Juicer was used with the 3D-DNA pipeline (v180922) (Dudchenko et al., 2017) with the following parameters ’-r 1-m haploid --build-gapped-map --sort-output’. The first round HiC and assembly files were manually curated locally within the Juicebox visualisation software (v1.11.08 for Windows) (Durand et al., 2016) the revised assembly file was resubmitted to the 3D-DNA post review pipeline with the following parameters ’--build-gapped-map --sort-output’ for final assembly and fasta files.

### Post-scaffolding assembly curation

Following Hi-C scaffolding, chromosome ends were assessed with Telociraptor (v0.9.0) (https://github.com/slimsuite/telociraptor) to correct for telomere-masking inversions or to remove contigs scaffolded onto telomeres. Five chromosome ends in hap1 and seven in hap2 were corrected to reveal telomeres. ChromSyn (Edwards et al., 2022) analysis revealed two hap2 chromosomes with arms mis-scaffolded end-to-end, which were also corrected with Telociraptor. ChromSyn and Telociraptor were also used to pair and rename hap1 and hap2 chromosomes. Each haploid assembly was then polished with Hypo (Kundu et al., 2019) using the partitioned PacBio and Illumina reads mapped onto the diploid assembly. Finally, Diploidocus was run using reads mapped onto the diploid assembly to identify and remove any haplotype-specific contigs that had been falsely duplicated, or unplaced “debris” contigs that were redundant following HyPo gap-filling.

### Genome quality assessment

We estimated the functional completeness of the genome using BUSCO (v5.1.2) (Simão et al., 2015a) in genome mode with the database, basidiomycota_odb10, and Metaeuk (Levy Karin et al., 2020) gene predictions. Basic genome statistics were determined with Quast (v5.0.2) (Gurevich et al., 2013). Kmer-based genome sequence completeness and accuracy was estimated with Merqury (v1.3) (Rhie et al., 2020).

### Mitochondrial genome assembly

We previously assembled and made public the complete circularized mitochondrial sequence (Tobias et al. 2022). We included the assembled Hv178a_MTDNA *Hemileia*

*vastatrix* isolate=Hv178a as a final contig within the principal pseudo-haplotype lodged at NCBI: JAMZDC000000000.

### Read depth genome size prediction and copy number analysis

Genome size predictions were made with DepthSizer (v1.6.3) (Chen et al., 2022), using partitioned PacBio reads (see below) mapped with Minimap (v2.4) (Li, 2018) and BUSCO (v5.1.2) (Simão et al., 2015a) gene predictions (see above). Read depth statistics and predicted copy numbers were generated using DepthKopy (v1.0.3) (Chen et al., 2022).

### Repeat family annotation

Transposable elements and simple sequence repeats were annotated with Repeat Modeler (v2.0.1) (Smit and Hubley, 2019) and Repeat Masker (v4.0.6) (Smit et al., 2019). Initial repeat libraries were constructed for the complete haplotype A genome and then used to predict repeats within both haplotype A and B with Repeat Maker using the following parameters; - pa 24 -gff -xsmall –lib. The softmasked output genome was used to map RNAseq data and for downstream annotation. We provide the Hv178a repeat data gff files here: https://osf.io/wtn4h/files/osfstorage.

### Protein annotation with RNA data from a closely related isolate

To annotate predicted coding genes, we mapped RNAseq data from the ancestral isolate, Hv178, to be used as evidence. First, we combined the Illumina paired-end RNAseq data derived from three replicates of the isolate Hv178 (v_2,3,5_) during compatible interaction with the respective differential host plant (*C. arabica* accession CIFC 34/13, S_H_2,3,5) at 3 key stages of infection, namely at 1, 4 and 10 days post inoculation. The raw Illumina data was first trimmed of adaptors and low-quality reads using default settings with fastp (v0.19.6) (Chen et al., 2018). We then mapped the data to the masked genome of haplotype A and B independently using hisat2 (v2.1.0) with ‘--no-unal’ and all other defaults. The resulting bam files were incorporated as evidence for annotation with Funannotate (v1.8.15) (Palmer and Stajich, 2023) with the predict function and using the BUSCO dikarya_odb9 database (Simão et al., 2015a) with diamond (v2.1.6) (Buchfink et al., 2021). The predicted amino acid (aa) fasta files for each haplotype were then put through Interproscan (v5.52-86.0) (Jones et al., 2014) to determine putative functions based on the following databases Pfam, COILS, Gene3D. The same aa fasta files were screened with SignalP (v4.1f) (Nielsen, 2017) and the subsequent 797 haplotype A and 891 haplotype B proteins with EffectorP (v3.0) (Sperschneider and Dodds, 2022) to identify predicted effectors.

### Mapping of Illumina data to Hv178a to investigate copy number variation

We investigated copy number (CN) of conserved single copy genes using Illumina data Hv178a and from 11 other *H. vastatrix* strains; Hv23, Hv70, Hv71, Hv178, Hv264a, Hv741, Hv995, Hv999, Hv1427, Hv3305 and Hv race I (Ángel C. et al., 2023a). For each strain, paired-end read data was mapped onto Hv178a haplotype A using BWA-Mem (Li, 2013). The CN for each gene in each strain was estimated using DepthKopy (Chen et al., 2022), based on empirical single-copy read depth estimates derived from single-copy BUSCO genes. The Copy Number Ratio (CNR) for each gene was then calculated as the ratio of the CN for Hv178a versus the mean CN for all other strains. The data was subject to a log2 transformation, where zero represents no CN difference, and a value of +/- 1 represents a two-fold difference.

### Copy number estimation by quantitative real-time PCR (qPCR)

#### Primer design, insert amplification and cloning

Validation of genomic CN prediction in chr17 was tested with qPCR analysis in isolate Hv178a compared with Hv178. Three gene targets were used: a predicted effector located in chr17A (Hv178aA_012933-T1), a conserved single copy gene located in chr 17A (putative fluoride ion transporter CrcB – locus 13767at5204) and a conserved single copy gene located in chr 6A (Actin-related protein 2-Mar complex subunit 5 – locus 82138at5204), denoted as E33, B17.1 and BC6, respectively. BUSCO conserved gene loci are available as supplementary data (https://osf.io/wtn4h/files/osfstorage). Four biological replicas of DNA extracted from isolates Hv178a and Hv178, as previously described (Silva et al., 2018), were used. Primers were designed using Primer Select (DNASTAR Lasergene, v 11.1.0.54). Amplification of the E33, B17.1 and BC6 gene fragments to use as cloning inserts was performed using DNA template from isolate Hv178a under the following conditions: 20 μL reactions containing 4 μL of GoTaq® Flexi buffer (1X, Promega, Madison, WI, USA), 2 μL of MgCl_2_ (25 mM, Promega, Madison, WI, USA), 0.5 μL of dNTPs mix (10 μM, Thermo Fisher Scientific, Waltham, MA, USA), 0.5 μL of BSA (10 mg/mL, GeneOn, Germany), 1 μL of each forward and reverse primers (10 mM, StabVida, Caparica, Portugal), 0.2 μL of GoTaq® Flexi DNA Polymerase (5 u/μL, Promega, Madison, WI, USA), 3 μL of DNA template (15 ng/ μL) and water to the final volume. A negative control with no template was included. PCR was carried out with an initial denaturation step at 94°C for 3 min, followed by 35 cycles of denaturation at 94°C for 1 min, annealing at the respective temperature for each gene (Table 1) for 1 min and extension at 72°C for 1 min, and one final step of extension at 72°C for 7 min. PCR products were purified using the Wizard® SV Gel and PCR Clean-Up System kit (Promega, Madison, WI, USA), quantified in a Qubit 4 Fluorometer (ThermoFisher Scientific, Waltham, MA, USA) and cloned into the vector and *E. coli* JM109 competent cells using the pGEM-T Easy Vector System II Systems Kit according to the manufacturer’s instructions (Promega, Madison, WI, USA), in a 3:1 (insert:vector) molar ratio. Positive colonies were confirmed by colony PCR using the universal primers SP6 and T7 and the specific primers for the inserts (E33, B17.1, BC6, Table 1). The cloned plasmids were isolated from pelleted cell cultures using Wizard® *Plus* SV Minipreps DNA Purification kit (Promega, Madison, WI, USA) and sequenced (Stabvida, Caparica, Portugal) to verify the presence and the correct insertion of the fragments in the vectors.

**Table 1.**
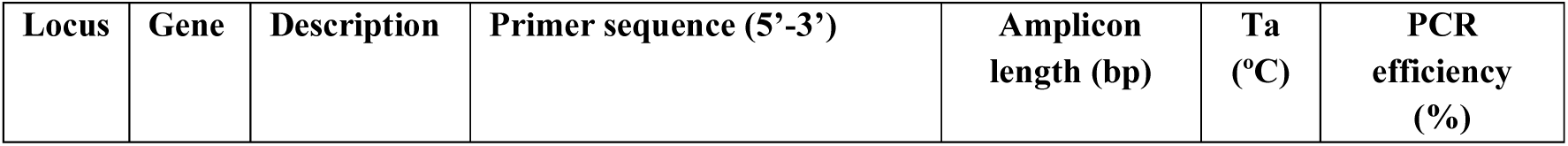
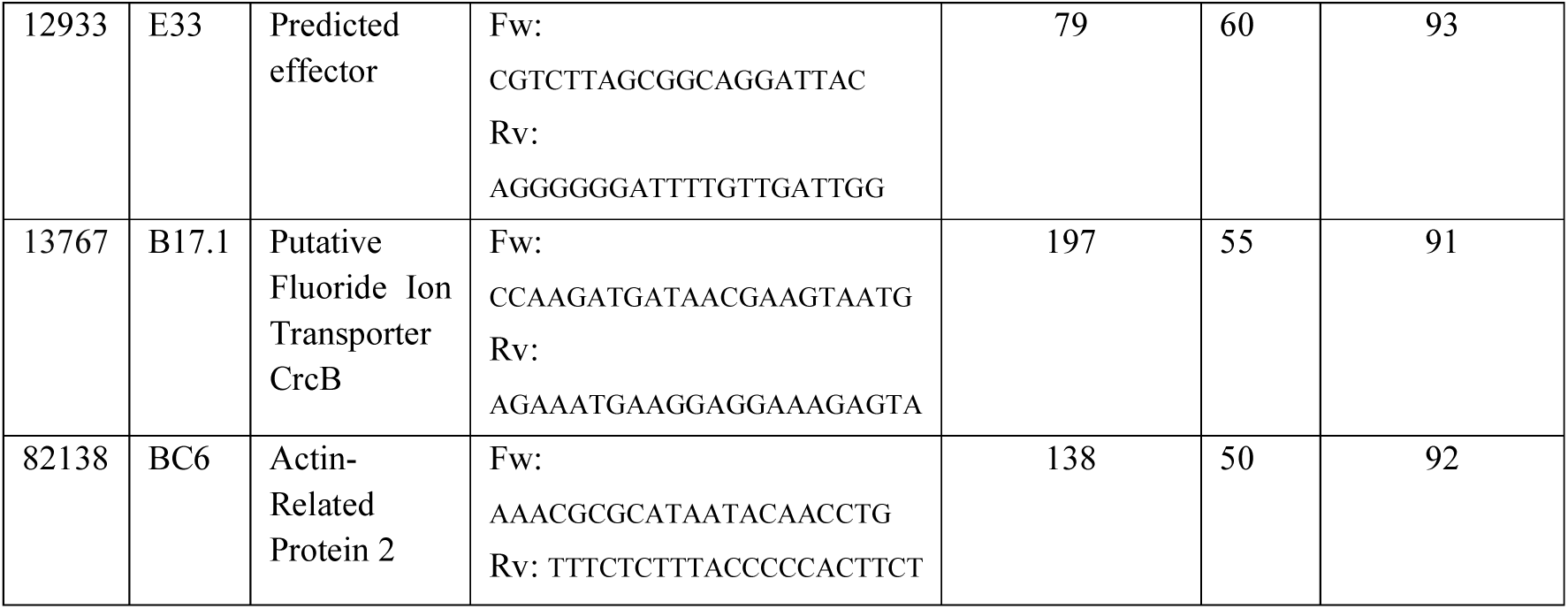
Primer sequences for qPCR analysis of target genes.

#### Construction of standard curves for determination of gene copy numbers

Plasmid DNA samples extracted from cell cultures cloned with E33, B17.1 and BC6 gene fragments were quantified using Qubit™ 1X dsDNA BR Assay Kit (Thermo Fisher Scientific, Waltham, MA, USA) and first diluted serially to 1 ng/µL and 0.01 ng/µL. The corresponding copy number of plasmid DNA per microliter was calculated using the following equation (Whelan et al., 2003) :

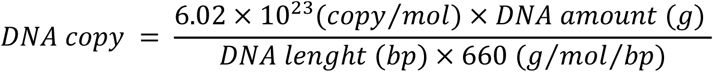

From a dilution of 100 copies/µL (0.01 ng/µL), subsequent dilutions were prepared for each target gene, corresponding to 75, 50, 25, 20, 15, 10, 5, 3, 2, 1.5, and 1 copy/µL. Dilutions from 100 to 1 plasmid copy/µL were run on a CFX Connect Real-Time PCR Detection System (Bio-Rad, Hercules, CA, USA) with the following conditions: 20 µL of reaction mixture prepared with 10 µL of SsoFast EvaGreen® Supermix (Bio-Rad, Hercules, CA, USA), 0.8 µL of each forward and reverse primers (0.01 mM), 1 µL of DNA template and water to the final volume. All qPCR experiments were carried out with the following thermal cycling protocol: an initial denaturation step at 95°C for 3 min, followed by 41 cycles of denaturation at 95°C for 15 s, annealing and extension at Ta (Table 1) for 30 s. In each set of reactions, at least 3 replicates per copy dilution were run, in addition to a non-template control. A standard curve for each gene was generated by plotting quantification cycle (Cq) average values (with standard deviation < 0.5) of each dilution step against the corresponding log10 transformed number of gene copies in the standard. Amplification efficiencies (E) were determined using LinRegPCR software (v 2013.0). All standard curves, generated by linear regression analysis of the plotted points, yielded strong linearity with correlation coefficients (R^2^) of ≥0.998 (Suppl. Fig. 1). Combined with amplification efficiencies over 0.90, these parameters qualify the standards for reliable gene copy number quantification (Svec et al., 2015).

#### Absolute quantification of gene copy number in qPCR

To determine the optimal sample concentration, providing a Cq value falling within the standard curve range for each gene, DNA pools of isolates Hv178a, and Hv178 were prepared by combining 1 ng/µL from 2 biological replicates per isolate, followed by serial dilution to obtain the following concentrations: 1 ng/µL, 0.1 ng/µL, 0.01 ng/µL, and 0.001 ng/µL. Each DNA pool was run in triplicate with the primers E33, BC6, and B17.1 as previously described. The amplification efficiency for each gene was determined using LinRegPCR (v2013.0). Henceforth, qPCR runs of 4 biological replicate samples from isolates Hv178a and Hv178 at 0.1 ng/µL, with 3 technical replicates for each sample, were performed with the primers E33, BC6, and B17.1 and analyzed as previously described. Dissociation curves (Suppl. Fig. 2) and agarose gel electrophoresis were used to assess the presence of non-specific PCR products. Mean Cq values from replicates (standard deviation ≤0.5) were used to calculate the copy number of each gene based on the corresponding standard curves. For each isolate, 3 biological replicates were selected for subsequent calculations. Gene copy number ratios were calculated between Hv178a and Hv178 for each target gene. To normalize gene copy number values considering the genome size of the two isolates, two alternative approaches were employed to estimate the number of genome copies present in the DNA template. In the first approach, genome copy numbers were determined by relating the amount of DNA template to the genome size of each isolate, as described by the following formula:

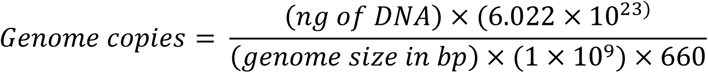

Genome sizes (Hv178a – 772Mbp; Hv178 – 774Mbp) previously determined by flow cytometry were used (Laureano, 2024). In the second approach, calculations were made using the mass (ng) equivalent to the two haploid nuclei of each isolate (Doležel et al., 2007), as follows:

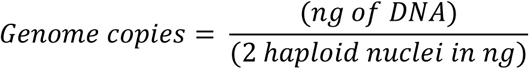

The calculation of the mass of the two haploid nuclei was based on the size of each isolate’s genome and the value of 1C (picograms of DNA in a haploid nucleus), using the conversion factor 1pg = 978 Mbp (Doležel and Bartoš, 2005). Absolute copy number was then divided by the genome copies retrieved from each formula, and the copy number ratios between Hv178a and Hv178 were calculated.

## Results and Discussion

### A complete pseudo-phased genome for *Hemileia vastatrix* (CIFC Hv178a)

We have assembled and pseudo-phased the di-karyotic genome for *Hemileia vastatrix* (CIFC Hv178a) using 200 X coverage with PacBio Sequel continuous long reads (CLR) followed by scaffolding chromosomes with chromatin conformation reads. Our genomes assembled to ∼660 Mbp, smaller than previously determined with flow cytometry at 796 Mb (Tavares et al., 2014). The most recently published *H. vastatrix* genome appears to confirm the 1C size at 775 Mbp (Ángel C. et al., 2023), however this genome is presented as a collapsed haploid assembly. It is likely that our smaller genome size is due to the earlier generation sequence technology data, known to be more error prone, and possible collapsing of highly repetitive regions during the assembly, polishing and scaffolding process. Indeed, our analysis predicted a genome size of 751.6 Mbp (hap A) to 767.2 Mbp (hap B), with an estimated empirical single copy sequencing depth of 157 X (data not shown), though our final assemblies are smaller. Conversely, it is also possible that the genome size reported by Ángel C. et al. (2023) is inflated due to the inclusion of haplotype-variable contigs within the collapsed assembly. Despite the size discrepancy our Hv178a genomes are highly complete with Benchmarking Universal Single-Copy Ortholog (BUSCO) (Manni et al., 2021; Simão et al., 2015) results of 82 and 76 percent (Table 2).

**Table 2.**
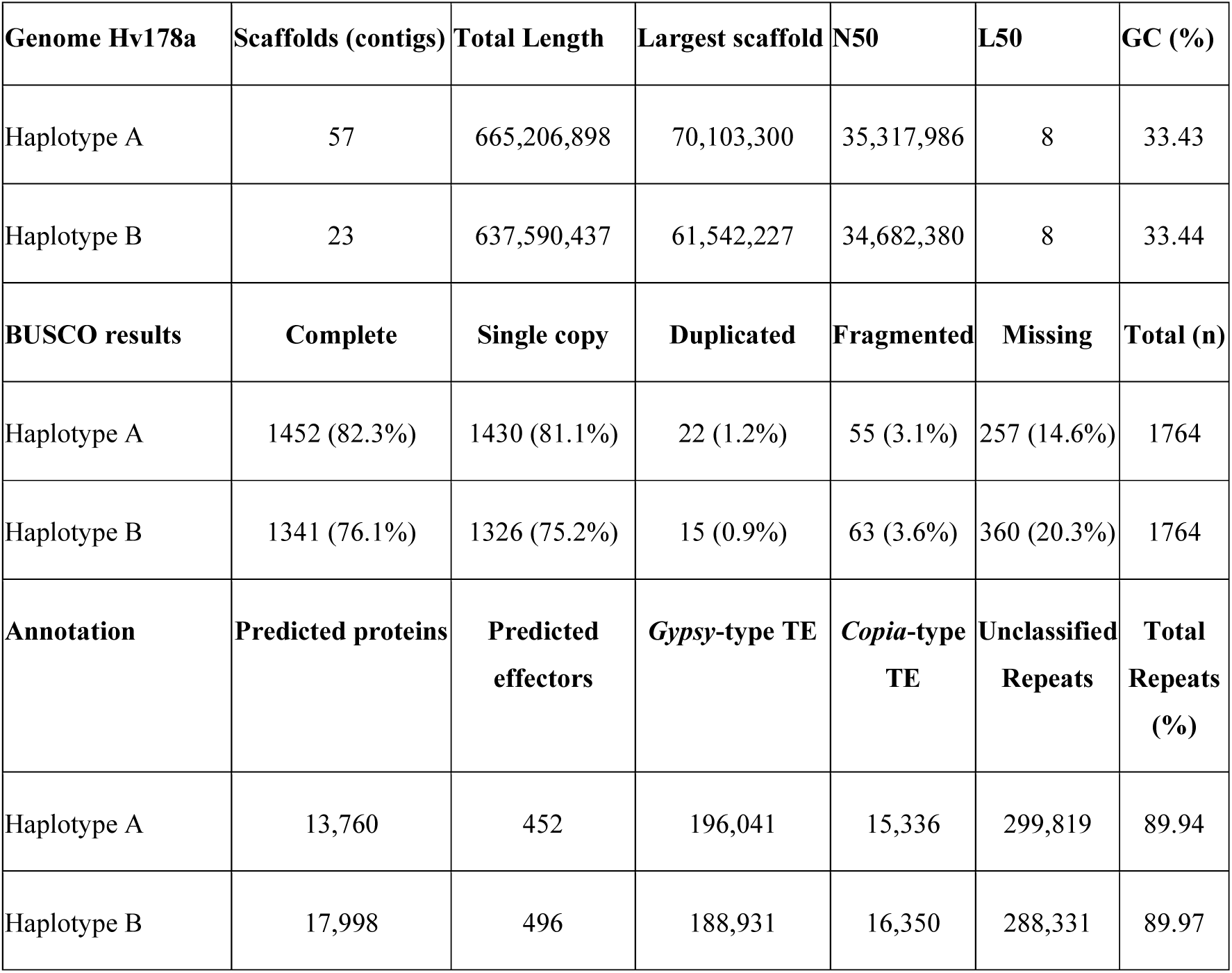
*Hemileia vastatrix* (strain CIFC Hv178a) HiC scaffolded genome assembly statistics using Quast *(Gurevich et al., 2013)* and BUSCO v. 5.1.2 *(Simão et al., 2015)* with basidiomycete_odb10 database.

### *Hemileia vastatrix* has 18 haploid chromosomes

The final Hv178a assembly has 18 haploid pseudochromosomes, clearly represented in the heatmaps (Figure 2), a number that is consistent with other rust fungi within the Pucciniamycotina subdivision (Schwessinger et al. 2020; Duan et al. 2022; Edwards et al. 2022). Until now, chromosome numbers have not been clearly assigned to this evolutionary group of rust-forming basidiomycetes. An earlier cytological study reported the difficulty in determining a conclusive number of chromosomes for *H. vastatrix* and suggested as the best resolved data the presence of 14 chromosomes (Chinnappa and Sreenivasan, 1965). More recently, Tavares et al., (2013) determined between 7-13 chromosomes using both electrophoretic and cytological karyotyping, as well as fluorescence in-situ hybridization of two races of *H. vastatrix* (race II - isolate 1065 and race VI - isolate 71) from the CIFC/ISA collection. However, despite being an ancestral rust from the family Mikronegeriaceae, chromosome numbers appear to be highly conserved. Genome data for Hv race I (Ángel C. et al., 2023) produced eleven telomere-to-telomere and seven near complete chromosomes, suggesting that further curation would align with the numbers that we determined for Hv178a.

**Figure 2.**
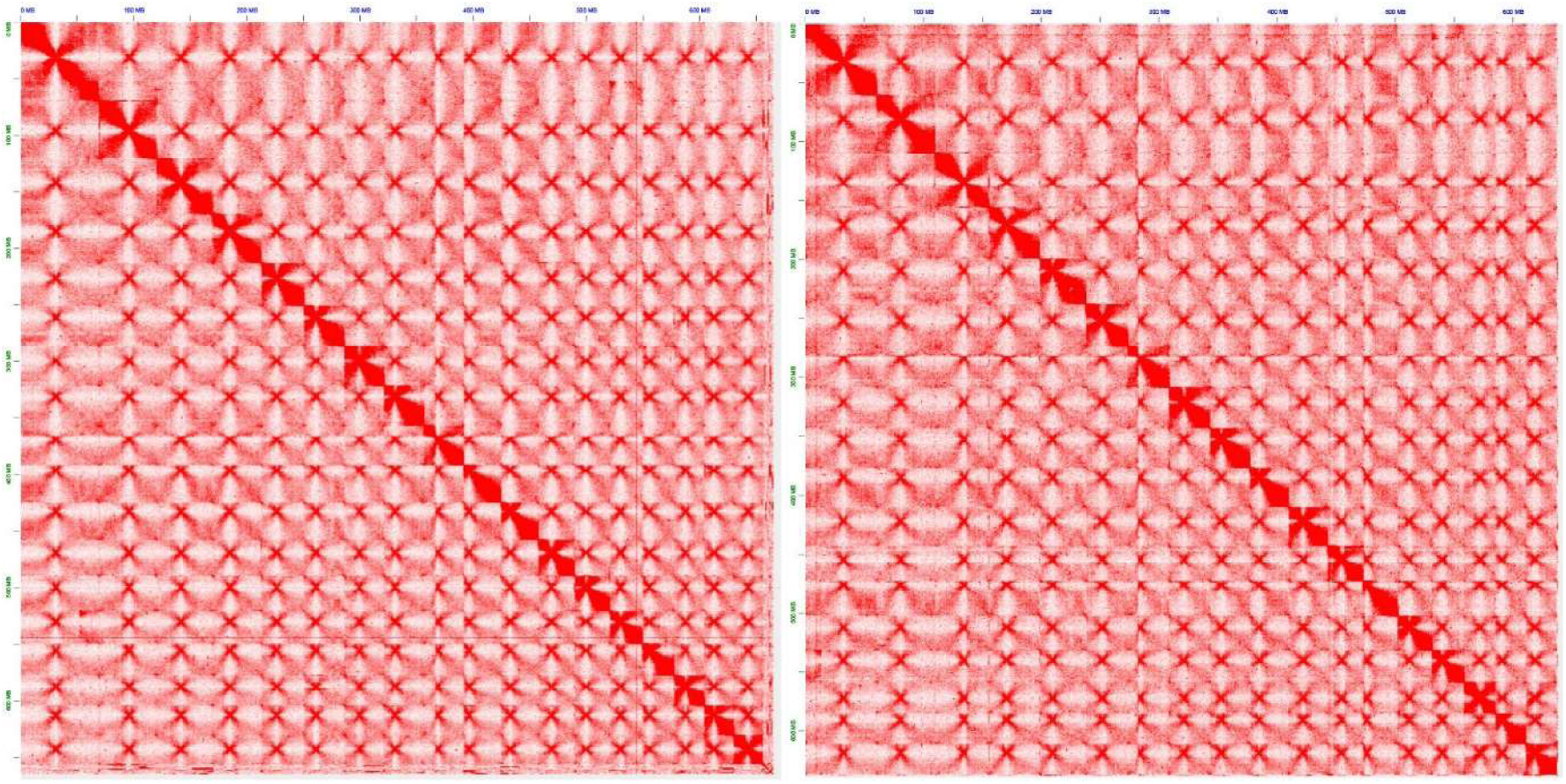
Heatmap image from Juicebox visualisation software (v 1.11.08) of the scaffolded *Hemileia vastatrix* (strain CIFC Hv178a) genome assembly shows 18 chromosomes visible in pseudo-haplotype A (left) and B (right).

### Repeat annotation reveals extensive *Gypsy*-type retrotransposons

The Hv178a di-haploid genomes were found to be extremely repetitive at 89.94% and 89.97% for haplotype A and B respectively, with low GC content at 33.44%. Similarly, Ángel C. et al (2023) determined 81.66% masked repeat sequences within Hv race I. On investigation of the class of repeat elements, we determined 196,041 and 188,931 *Gypsy*-type long terminal repeat retrotransposons (LTR) as well as 15,336 and 16,350 *Copia-*type LTRs. In all, we determined 54.56% LTR retrotransposons and ∼34.50% unclassified repeats for each haplotype. Based on Cristancho et al (2014) and Porto et al. (2019) genome assemblies, Orozco-Arias et al.(2022) improved the annotation of transposable elements (TEs) in *H. vastatrix* showing that lineages from the Gypsy superfamily correspond to 93.3% of all the LTR-RTs identified. The expansion of transposable elements (TE) classified within the *H. vastatrix* genome is comparable to other large rust fungal genomes such as *Austropuccinia psidii* (Tobias et al., 2021), *Phakopsora pachyrhizi* (Gupta et al., 2023) and *Puccinia polysora* (Liang et al., 2023). According to Orozco-Arias et al. (2022), the recent activity of three Gypsy lineages (CO-HUI, Soroa, and V_Clade) might be at the origin of the genome size expansion of *H. vastatrix*. Expansion of transposable elements has been suggested to play a role in host range adaptation (Gupta et al., 2023), and the low GC content may be a result of DNA methylation silencing of TEs leading to deamination of methylated cytosines over time, as proposed for *A. psidii* (Tobias et al., 2021). In *P. pachyrhizi* (Gupta et al., 2023), the patterns of expression of TEs were studied during the infection cycle and a subset determined to be highly expressed *in planta* at 24 hours post inoculation. It would be interesting to determine if a similar pattern of gene expression is observed during infection on *C. arabica*, particularly on differential host plants, as a potential explanation for new virulence mechanisms.

### Predicted protein coding gene numbers are different in each pseudo-haplotype

We annotated the predicted protein coding genes within each haplotype and determined 13,760 recognized as a common feature of dikaryotic rust fungi, where the two haploid nuclei that coexist within the same cell can evolve semi-independently over long evolutionary periods. High-quality phased genome assemblies of rust fungi were able to demonstrate that the two haplotypes may display substantial sequence divergence as well as structural variation, gene presence–absence polymorphisms, and haplotype-specific genes (Luo et al., 2025; Wang et al., 2024).The difference in numbers of annotated genes between haplotypes in *H. vastatrix* could be difficult to understand given the lack of any known sexual stage, albeit population genomic footprints of hybridization and recombination were previously detected (Silva et al., 2018). However, long-term asexual reproduction can lead to divergent evolution within the two dikaryotic nuclei by promoting to a continuous accumulation of mutations and TEs, structural variations and a high level of heterozygosity as shown in asexual lineages of *P. striiformis* f.sp*. tritici* isolates (Schwessinger et al., 2020). We processed the amino acid sequences to predict subcellular localization based on known signal peptides (Nielsen, 2017). Based on the outputs of mature proteins, that had no recognisable transmembrane signal peptide, we predicted 452 and 496 virulence molecules, known as effectors in pseudo-haplotypes A and B. Effectors are characterised as secreted proteins that facilitate infection and suppress host defences (Sperschneider and Dodds, 2022). The predicted effectors were classified as either cytoplasmic or apoplastic based on machine learning approaches. This method identified 351 (406) cytoplasmic and 101 (90) apoplastic effectors in pseudo-haplotypes A and B. Based on 15 obligate biotroph genomes tested, average percentages of cytoplasmic and apoplastic predicted effectors are 32.5 and 11.3% respectively (Sperschneider and Dodds, 2022). In the haploid genomes for Hv178a the percentages of cytoplasmic and apoplastic predicted effectors are 44.0 and 12.7% (haplotype A), 45.6 and 10.1 (haplotype B) which is more comparable to the finding of 40% apoplastic effectors determined for *A. psidii*, a similarly larger genome for an obligate biotroph (Tobias et al., 2021). Using ChromoMap (Anand and Rodriguez Lopez, 2022) we show the predicted effector locations and numbers are different across chromosome pairs indicating that the two nuclei likely do not interact and are on their own evolutionary path (Figure 3). The differences observed here suggest that the two nuclei of *H. vastatrix* may harbor partially distinct virulence repertoires, potentially increasing the adaptive potential of this obligate biotrophic pathogen.

**Figure 3.**
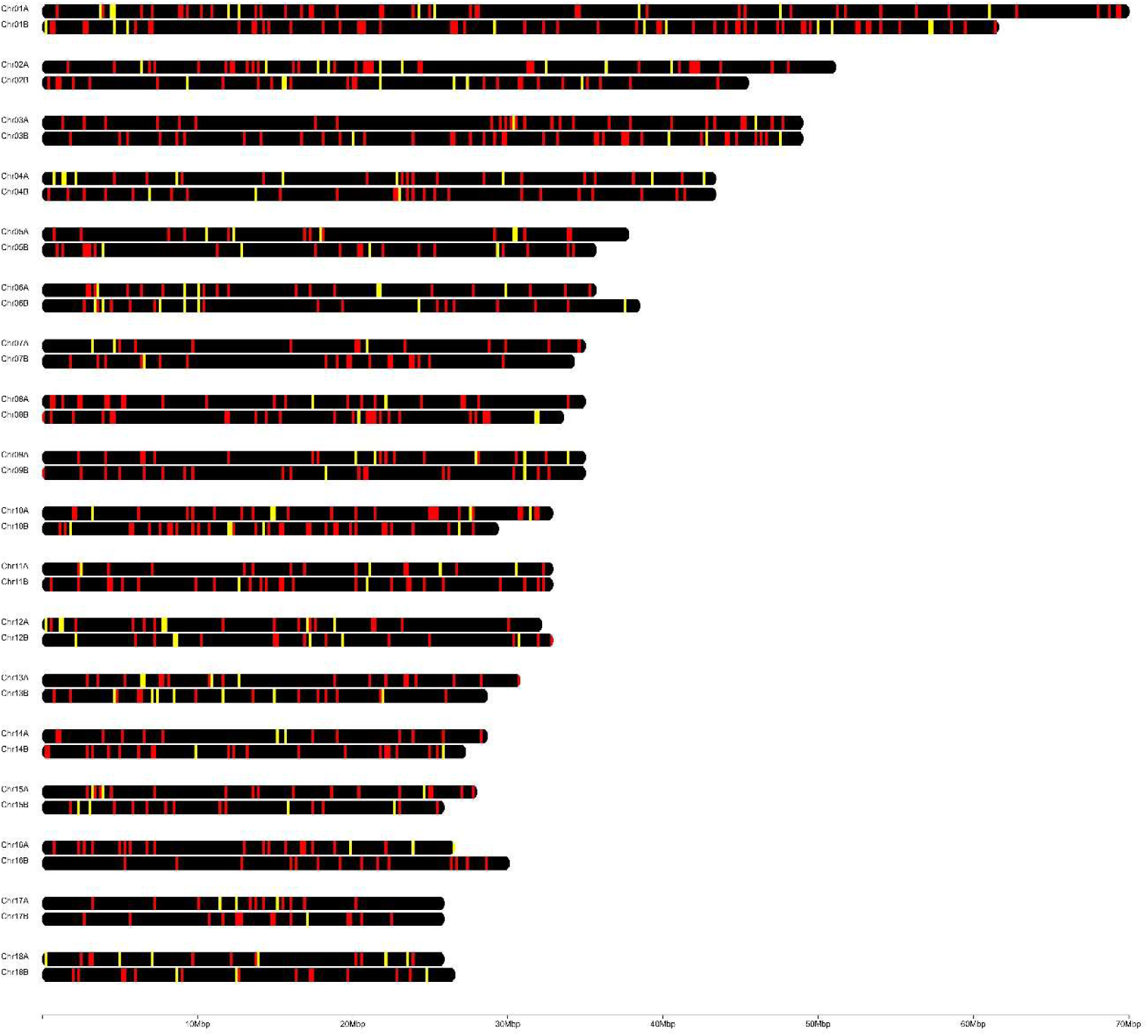
Predicted cytoplasmic (red) and apoplastic (yellow) effector gene locations on the pseudo-phased chromosome pairs for CIFC *Hemiliea vastatrix* 178a visualized using ChromoMap (Anand and Rodriguez Lopez, 2022). Lines represent single predicted genes.

### Read depth analysis suggests Chr17 trisomy in Hv178a

To investigate whether any genes had gained or lost copy number in Hv178a, the Copy Number Ratio (CNR) of Hv178a compared to eleven other *H. vastatrix* strains (Hv23, Hv70, Hv71, Hv178, Hv264a, Hv741, Hv995, Hv999, Hv1427, Hv3305 and HvI) was calculated for each gene (Suppl. Data 1). To identify potential regions of interest, log2 CNR was plotted against chromosomal position for each chromosome (Suppl. Fig. 3). For most chromosomes, such as Chromosome 1, log2 CNR was around 0 across the length of the chromosome (Fig 4a). However, Chromosome 17 displayed a markedly different pattern, with a consistent elevation of 0.5 or more (Fig 4b). This corresponded to a CNR value of 1.4-1.5. Given the noisiness of the CN estimates, which were based on short-read data, rather than more consistently sequenced long-reads, this was highly suggestive of a 50% increase across the entire chromosome, which could be explained by a third copy (trisomy) of Chromosome 17 in the genome.

**Figure 4.**
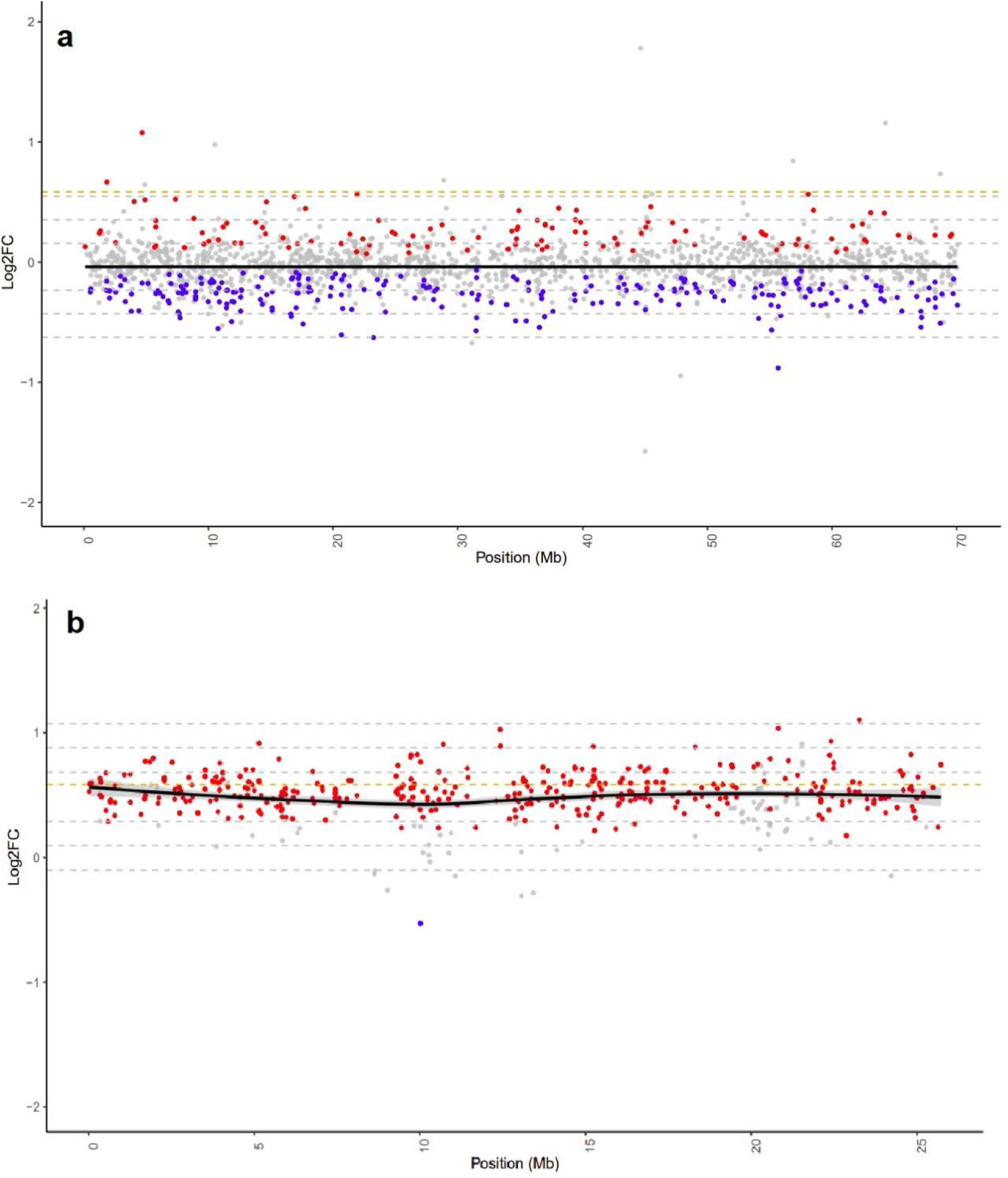
Hv178a Log2 Copy Number Ratio compared to the mean of 11 other Hv strains. a) Chromosome 1, b) Chromosome 17. The CN ratio was calculated as the CN for Hv178a versus the mean of 11 other *H. vastatrix* strains: Hv23, Hv70, Hv71, Hv178, Hv264a, Hv741, Hv995, Hv999, Hv1427, Hv3305 and HvI. A log2 CN ratio of 0 indicates no CN difference. Each dot is a gene, plotted against chromosomal position. Red, Hv178a has the highest CN of all strains; Blue, Hv178a had the lowest CN of all strains; Grey: Hv178a is not the most extreme CN. Black line, loess regression line of log2 CNR; Grey lines, 1-3 standard deviations from the mean. For all chromosome plots, see Supplementary Figure 3.

### Quantitative PCR supports Chr17 trisomy in isolate Hv178a

The putative trisomy of chromosome 17 in isolate Hv178a was further evaluated by qPCR absolute quantification of target gene copy number, using isolate Hv178 - the original field-collected strain from which the mutant Hv178a was produced- as the reference. Under the hypothesis of chromosome 17 trisomy, loci located on chromosome 17 are expected to be present in three copies in the trisomic isolate versus two copies in the euploid isolate, while loci on other chromosomes are expected to remain disomic in both isolates. Accordingly, when normalizing absolute copy numbers between isolates, a ratio of approximately 1.5 (3:2) should be observed for both chromosome 17 loci analyzed (effector gene E33 and BUSCO gene B17.1), whereas a ratio of approximately 1.0 (2:2) should be observed for the chromosome 6 BUSCO gene (BC6), serving as a disomic reference chromosome. Indeed, estimation of absolute gene copy number for these three gene targets showed ratios between Hv178a and Hv178 of 1.601 ± 0.311for E33, 1.220 ± 0.311 for BC6 and 1.472 ± 0.384 for B17.1. These proportions are maintained with very similar values when a genome size normalization factor is additionally applied (Table 2). Although a previous estimate of the genome size for Hv178a was reported by Tavares et al. (2014) as 796.8 Mbp, we used the values reported by Laureano (2024) for normalization purposes, 772 Mbp for Hv178a and 774 Mbp for Hv178. These estimates were selected because they were based on a larger number of replicates for Hv178a (n = 10 vs. n = 5 in Tavares et al., 2014) and because they provide the only available genome size estimate for Hv178. Nevertheless, we also calculated the absolute copy number normalized using the genome size reported by Tavares et al. (2014), and no significant differences in ratios were observed (E33-1.648 ± 0.320; BC6 - 1.256 ± 0.331; B17.1 - 1.515 ± 0.395). This consistent observation of a ∼1.5-fold increase in absolute copy number for chromosome 17 loci, coupled with a ∼1.0 ratio for the chromosome 6 control locus, provides quantitative evidence supporting trisomy of chromosome 17 in Hv178a (Suppl. Table 1). This dual-locus strategy on the same chromosome, combined with an independent chromosomal control, minimizes locus-specific amplification bias and strengthens the inference of chromosomal copy number variation.

Collectively, these findings are consistent with a third copy of chromosome 17 in Hv178a, which could mean that the acquisition of increased virulence by an additional virulence factor (v4) in relation to Hv178 was accompanied by a catastrophic genomic event. Such extensive rearrangements may be more common in coffee rust than previously anticipated, as other mutant isolates derived from adaptation experiments made at CIFC exhibit substantial genome size differences, for example, Hv264 (v_1,4_) to Hv264a (v_1,4,6_) show an increase of 36 Mbp (Laureano, 2024). These results suggest that host adaptation may act as catalyst for large-scale genomic restructuring.

Structural and numerical genomic variations are increasingly recognized as important contributors to rapid adaptive evolution in fungal plant pathogens. Supernumerary or accessory chromosomes have been well-studied in *Fusarium* species (Coleman et al., 2009; Ma et al., 2010; Vanheule et al., 2016) and *Zymoseptoria tritici* (Croll et al., 2013; Goodwin et al., 2011), and typically are enriched in effector genes and transposable elements, offering distinct structural and functional compartmentalization within a genome. As in the dispensable accessory chromosomes of *Fusarium* spp, mobile mini-chromosomes in the rust pathogen *Magnaporthe oryzae* are lineage-specific but show in addition extensive recombination with core chromosomes and *in planta* specific effector gene expression (Ma and Xu, 2019; Peng et al., 2019). These mini chromosomes likely serve as repositories of virulence genes that can be gained, lost, or rearranged at higher frequencies than core genomic regions, promoting rapid shifts in the pathogen’s interaction with host resistance. The extra chr 17 chromosomal copy in Hv178a may similarly expand the genomic landscape available for effector gene diversification and expression, analogous to how mini-chromosomes in *M. oryzae* function as fast-evolving genomic compartments driving host adaptation. Genomic regions with elevated structural plasticity are hotspots for generating virulence-associated variation, enabling rapid responses to host immune pressure. Such change could potentially increase the copy number, dosage imbalance and combinatorial function of effector loci, enhancing the pathogen ability to evade host recognition or suppress immunity, contributing to increased virulence.

## Data availability

Raw data is available at the following BioProject accession number at the National Centre for Biotechnology Information (NCBI): PRJNA837996. The genomes have been deposited under the accessions PRJNA851216, JAMZDC000000000 and PRJNA851215, JAMZDD000000000. Protein annotation files, including predicted effector data, are available at: https://osf.io/wtn4h/files/osfstorage. Isolate data used for annotation and for copy number analysis are available at the request of the authors.

## Acknowledgements

We thank Ana Paula Pereira and the technical staff from CIFC/ISA, namely Célia Lopes, Idalina Gomes and Miguel Ribeiro, for the support provided on isolate multiplication and pathotype testing, as well as for the maintenance of coffee plants and preservation of CIFC rust collection.

## Author contributions

PAT assembled nuclear and mitochondrial genome assemblies, scaffolded the genomes with HiC reads, ran analyses, wrote much of the manuscript. RJE extensively curated the genome data, ran analyses, prepared figures, contributed to the methods and results of the manuscript writing. JB ran CN analyses and prepared figures. GL ran qPCR analyses and prepared figures. ID designed qPCR primers and participated in analyses. DB and VI extracted and purified HMW DNA. LGG prepared crosslinked spore samples for Hi-C sequencing. VV and MCS provided spore samples from CIFĆs collection, coordinated isolate multiplication and characterization, and contributed to the manuscript. RFP participated in sequencing funding and contributed to the manuscript. DB initiated and led the research, sourced grant funding, prepared samples for PacBio, for Illumina and for Hi-C sequencing and wrote much of the manuscript. All authors contributed to and read the manuscript.

