## Supplementary material for "A pseudo-phased genome assembly for *Hemileia vastatrix* reveals an isolate-specific chromosomal haploid trisomy": Suppl. Fig. 1

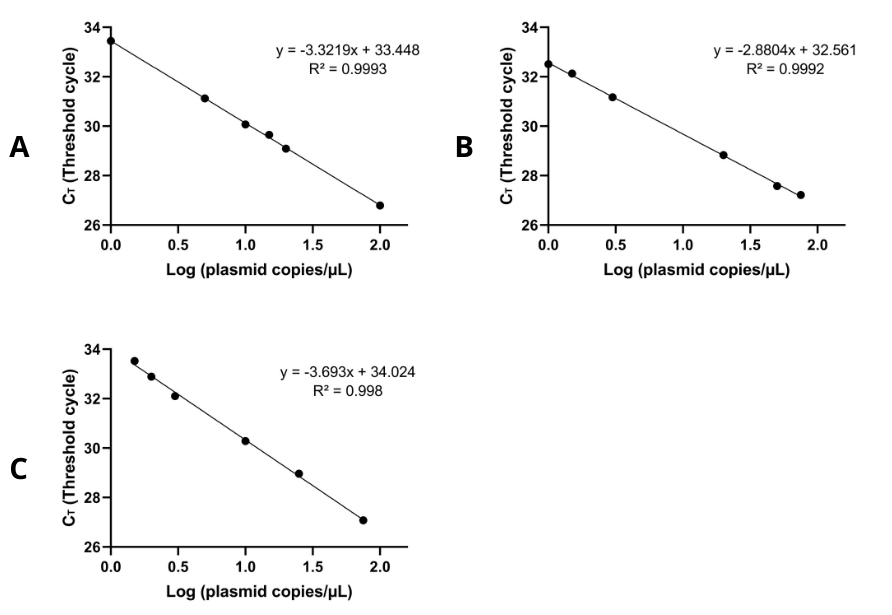

Supplementary Figure S1. Standard curves of the plasmid copy numbers for each target gene obtained by plotting the average Cq of 6 points against the logarithm of the number of copies in 1 µL for each sample: E33 (A); BC6 (B); B17.1 (C). Each curve presents its own equation and R^2^.
