## Supplementary material for "A pseudo-phased genome assembly for *Hemileia vastatrix* reveals an isolate-specific chromosomal haploid trisomy": Suppl. Fig. 2

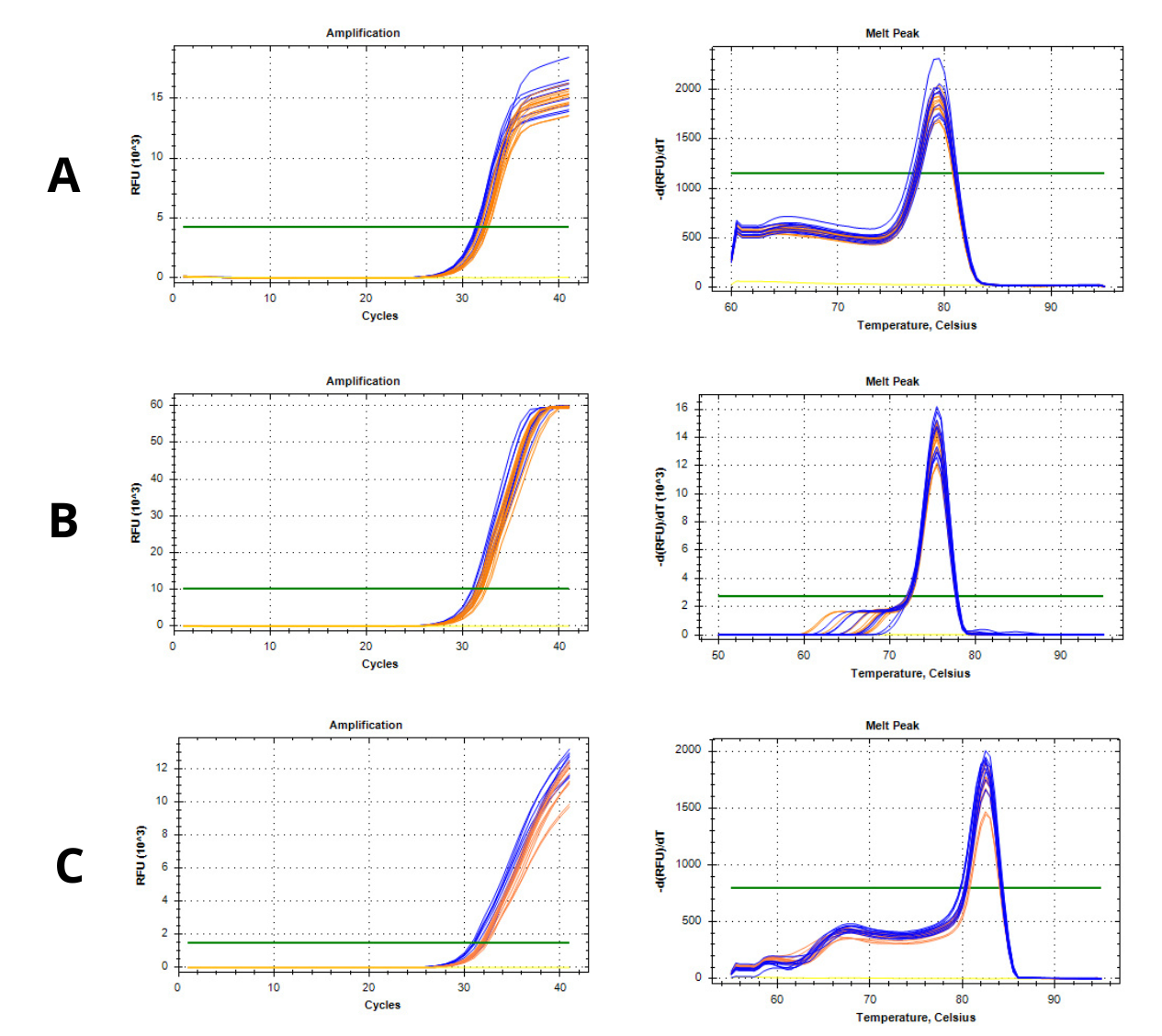

Supplementary Figure S2. Amplification curves and dissociation curves from qPCR analysis of Hv178a and Hv178 samples at 0.1 ng/µL for E33 (A), BC6 (B), and B17.1 (C) target genes. Samples of Hv178a are shown in blue, while samples of Hv178 are shown in orange. The negative control is shown in yellow. (Bio-Rad CFX Maestro 1.0 software, version 4.0.2325.0418, Bio-Rad, USA).
