## Supplementary material for "A pseudo-phased genome assembly for *Hemileia vastatrix* reveals an isolate-specific chromosomal haploid trisomy": Suppl. Table 1

|  | Absolute copies | | |  | Copies/genome size by nanogram | | |  | Copies/genome mass by nanogram | | |
| --- | --- | --- | --- | --- | --- | --- | --- | --- | --- | --- | --- |
|  | Hv178a | Hv178 | Ratio  Hv178a/Hv178 |  | Hv178a | Hv178 | Ratio  Hv178a/Hv178 |  | Hv178a | Hv178 | Ratio  Hv178a/Hv178 |
| E33 | 9.842 ± 2.899 | 6.081 ± 0.423 | 1.601 ± 0.311 |  | 0.833 ± 0.200 | 0.516 ± 0.029 | 1.597 ± 0.310 |  | 1.554 ± 0.374 | 0.962 ± 0.055 | 1.597 ± 0.310 |
| BC6 | 18.459 ± 6.440 | 15.033 ± 0.447 | 1.220 ± 0.311 |  | 1.562 ± 0.445 | 1.275 ± 0.031 | 1.217 ± 0.320 |  | 2.914 ± 0.830 | 2.379 ± 0.058 | 1.217 ± 0.320 |
| B17.1 | 17.402 ± 5.869 | 11.775 ± 0.333 | 1.472 ± 0.384 |  | 1.472 ± 0.405 | 0.999 ± 0.023 | 1.468 ± 0.383 |  | 2.747 ± 0.757 | 1.864 ± 0.043 | 1.468 ± 0.383 |

Suppl. Table 1 – Estimation of absolute copy number in isolates Hv178 and Hv178a, and derived ratios, for the gene targets studied using three different approaches: direct calculation from the standard curves (Absolute copies), normalization by the respective genome size of each isolate (Copies/genome size/ng), and normalization by the respective genome mass of each isolate based on the value of 1C in pg (Copies/genome mass/ng).
