## Supplementary figures and images for "A pseudo-phased genome assembly for *Hemileia vastatrix* reveals an isolate-specific chromosomal haploid trisomy"

### Suppl. Fig. 3

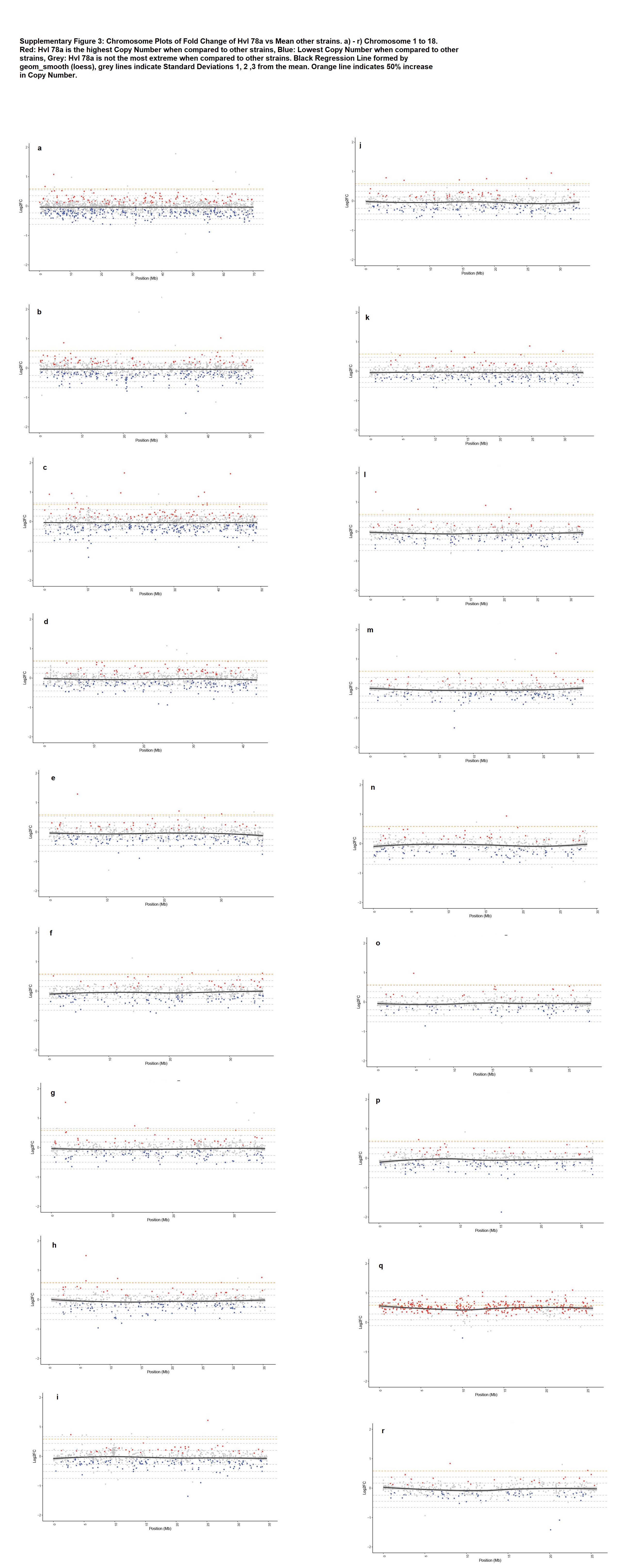
